# Sclerostin depletion induces inflammation in the bone marrow of mice

**DOI:** 10.1101/2020.11.01.364158

**Authors:** Cristine Donham, Betsabel Chicana, Alexander Robling, Asmaa Mohamed, Sonny Elizaldi, Michael Chi, Brian Freeman, Alberto Millan, Deepa Murguesh, Gabriela G. Loots, Jennifer O. Manilay

## Abstract

Romosozumab, a humanized monoclonal antibody specific for sclerostin, has been approved for treatment of post-menopausal women with osteoporosis at high risk for fracture. In several Phase III clinical trials, romosozumab decreased the risk of vertebral fractures up to 73% and increased total hip area bone mineral density by 3.2%. Previous work in 12 to 15-week-old sclerostin-knockout (*Sost*^-/-^) mice indicated that changes in immune cell development occur in the bone marrow (BM), which could be a possible side effect to follow in human patients. Our overall goal was to define the mechanisms that guide behavior of long-term hematopoietic stem cells (LT-HSCs) after exposure to an irregular BM microenvironment. SOST plays an important role in maintaining bone homeostasis, as demonstrated by the increased ratio of bone volume to total volume observed in *Sost*^-/-^ mice. Here, we examined the effects of short-term sclerostin depletion in the BM on hematopoiesis in young (8 week-old) mice receiving sclerostin-antibody (Scl-Ab) treatment for 6 weeks, and the effects of long-term Sost-deficiency on wild-type (WT) LT-HSCs transplanted into older (16-22 week-old) cohorts of *Sost*^-/-^ mice. Our analyses revealed an increased frequency of granulocytes and decreased frequency of lymphocytes in the BM of Scl-Ab treated mice and WT→*Sost*^-/-^ hematopoietic chimeras, indicating myeloid-biased differentiation in *Sost*-deficient BM microenvironments. This myeloid bias extended to extramedullary hematopoiesis in the spleen and was correlated with an increase in inflammatory cytokines TNFα, IL-1α and MCP-1 in the serum of the *Sost*^-/-^ BM. Additionally, we observed alterations in erythrocyte differentiation in the BM and spleen of *Sost*^-/-^ mice. Taken together, our current study indicates novel roles for *Sost* in the regulation of myelopoiesis and control of inflammation in the BM. Our animal studies strongly recommend tracking of hematopoietic function in patients treated with romosozumab.

## INTRODUCTION

Two rare human bone disorders, sclerosteosis and van Buchem disease, are characterized by dramatic increases in bone mineral density (BMD) and have been genetically traced to mutations in the *SOST* gene locus, which codes for sclerostin (1, 2). Sclerostin (*SOST*) is predominantly secreted by osteocytes and functions as a negative regulator of bone formation. Through the binding of low-density lipoprotein related protein 5 and 6 (LRP5 and LRP6), SOST has been shown to inhibit canonical Wnt signaling (3). LRP4 is also required for Wnt inhibition by sclerostin (4). Additionally, when Wnt is inhibited, osteoprotegerin (OPG) is decreased, and there is an increase in bone resorption (5). In *Sost*^-/-^ mice where Wnt is upregulated, OPG is concurrently increased (5). This demonstrates that *Sost* not only regulates bone formation, it also regulates bone resorption. The increase in BMD in *Sost*^-/-^ mice and in humans with decreased SOST function has led to consideration of sclerostin as a potential target for treatment of osteoporosis and other diseases associated with low bone mineral density. A humanized monoclonal antibody against sclerostin termed romosozumab (brand name Evinity) has been approved in the U.S. for women with osteoporosis after menopause at high risk for fracture. In several phase III clinical trials, romosozumab was shown to decreases the risk of vertebral fractures up to 73% and increase total hip area BMD by 3.2% (6).

Hematopoiesis occurs mainly in the bone marrow (BM). There is increasing evidence on the effects of altered bone homeostasis on hematopoiesis (7, 8). Self-renewal and quiescence of long-term hematopoietic stem cells (HSCs) and hematopoietic differentiation involves many cell signaling pathways, including Wnt, Notch, FGF and Hedgehog (9). Several studies regarding Wnt signaling and hematopoiesis have produced contradictory results (10-12). However, there have been a few reconciling propositions, including the idea that canonical Wnt signaling regulates hematopoiesis in a dosage-dependent fashion, Wnt signaling differentially affects fetal and adult HSCs, and interactions between canonical and non-canonical Wnt pathways influence the interpretation of these results (13).

Under homeostatic conditions, long-term hematopoietic stem cells (LT-HSCs) maintain a quiescent state in the BM, providing a lifelong reservoir of multipotent cells that replenish hematopoietic populations as they are depleted by use or age. Since the discovery of HSCs in 1961 (14), a growing list of hematopoietic stem and progenitor cell (HSPC) populations have been described, along with a set of cell surface markers to define them. Here, unless otherwise specified, we define the conglomerate HSPCs as LSKs (Lin^-^, Sca1^+^, cKit^+^), and LT-HSCs as LSK, CD150^+^, CD48^-^, Flk2^-^ (15-17). In an adult mouse, LT-HSCs give rise to all mature blood lineages. As cells progress from LT-HSCs to short-term hematopoietic stem cells (ST-HSCs) and then to multipotent progenitors 2, 3 or 4 (MPP2, MPP3, or MPP4), cells are generally thought to split into two different paths, the myeloid and lymphoid pathways. It has been demonstrated that MPP2 and MPP3 contribute largely to myeloid cells, and MPP4 to lymphoid cells (18). Lymphopoiesis produces cell types such as mature B, T and NK cells, whereas myelopoiesis produces a larger variety of mature cells, such as monocytes, granulocytes and erythrocytes.

Organismal aging is characterized by increased inflammation and decreased HSC functional capacity. This general decline in HSC function includes a reduction in long-term repopulating potential, homing and engraftment after transplantation (19), a decrease in lymphopoiesis and increase in myelopoiesis (20, 21) as well as an increase in total number of LT-HSCs (22). Aging is also associated with a chronic inflammatory phenotype that has been characterized by anemia, immunosenescence, thrombocytosis (23) as well as overproduction of inflammatory cytokines IL-1, TNF-α and IL-6 (24).

In 1978, the concept of a “hematopoietic stem cell niche” was introduced and is still being explored today (25). The spatial localization of HSC niches in the BM remains controversial, with early work suggesting that hematopoiesis is maintained in homogenously distributed niches, while more recent work suggests distinct niche structures (26). However, it is agreed upon that local BM microenvironments maintain HSCs and regulate their function by producing factors that act directly on HSCs such as SCF, CXCL12 and thrombopoietin. SCF binds to the KIT receptor on HSCs and is required for HSC maintenance (27). CXCL12 is involved in HSC maintenance and retention in the BM through activation signaling of CXCR4 on HSCs (28). CXCL12-CXCR4 signaling also regulates several myeloid and lymphoid progenitors’ proliferation and retention in the BM (29, 30). Thrombopoietin activates signaling through myeloproliferative leukemia protein on HSCs and is required for HSC maintenance (31). Thrombopoietin is also important in megakaryocyte and platelet production (32).

Our previous work demonstrated that *Sost*-deficiency in mice affects B lymphocyte development in a non-cell autonomous manner, CXCL12 levels are diminished in the bones of *Sost*^-/-^ mice, but that Wnt signaling in B lymphocytes does not appear to be affected (33). In this study, we extended our investigation to focus on HSC function and fate in sclerostin-deficient BM microenvironments. Specifically, we hypothesized that removal of sclerostin would influence LT-HSC self-renewal, maintenance and differentiation.

## MATERIALS AND METHODS

### Study Design/Statistical Analysis

G*Power 3.1 software was used to calculate the sample size required per group for all experiments. Specifically, the type of power analysis used was a priori: compute required sample size given α, power and effect size, where n1 = n2 (allocation ratio N2/N1 =1), α = 0.05, two tails, normal parent distribution and effect size of d =0.5. All box and whisker plots are composed of the ends of the box corresponding to the upper and lower quartiles, with the median marked by the vertical line inside. The whiskers correspond to the highest and lowest observations. All line data were expressed as the mean ± standard deviation. For flow cytometry results, statistical analysis was done using Mann-Whitney U-test or Student’s t-test with a two-tailed distribution, with two-sample equal variance (homoscedastic test) using GraphPad Prism. For all tests, p < 0.05 was considered to be statistically significant.

### Mice

C57BL/6 mice (Stock No: 000664) and B6.SJL (CD45.1, Stock No: 002014) were purchased from The Jackson Laboratory. *Sost*^*-/-*^ LacZ-knockin mice, in which *Sost* is globally deleted, have been previously described (33). Mice of both sexes were used on the C57BL/6 background and housed in sterile, microisolator cages with autoclaved feed and water. The UC Merced or Lawrence Livermore National Laboratories IACUCs approved all animal work.

### Flow cytometric analysis of bone marrow cells

Procedures for euthanasia, dissection and preparation of femurs and tibias were performed as previously described (33). Antibodies used are listed in Supplemental Table 1 and were purchased from eBioscience, BioLegend, Miltenyi Biotec, and BD Biosciences. A list of cell surface markers for the cell populations we examined is provided in Supplemental Table 3. Staining of all cells included a pre-incubation step with unconjugated anti-CD16/32 (clone 2.4G2 or clone 93) mAb to prevent nonspecific binding of mAbs to FcγR, except when staining for CD16/32. For extracellular staining, the cells were incubated with a panel of biotinylated mAbs for 20 minutes on ice, followed by a secondary stain with fluorochrome-conjugated streptavidin and additional primary directly conjugated mAbs for 20 minutes on ice and then 10 min at room temperature protected from light. DAPI or PI was used as a viability dye. Single color stains were used for setting compensations and gates were determined by historical data in addition to fluorescent-minus-one control stains. Flow cytometric data was acquired on the BD LSR II or BD Aria III. The data was analyzed using Flowjo, version 10 (Tree Star, Inc.).

### Scl-Ab treatment

SOST antibody, (Scl-Ab; Regeneron) (25 mg/kg), or vehicle (PBS) was administered subcutaneously to C57BL/6 mice starting at 8 weeks of age, twice weekly, for 6 weeks.

### uCT analysis

Formalin-fixed femora from vehicle- and Scl-Ab-treated mice were scanned, reconstructed, and analyzed as previously described (34). Briefly, 10-μm resolution, 50-kV peak tube potential and 151-ms integration time were used to collect scans on a Scanco uCT-35 tomographer. The distal 60% of each femur was scanned. Standard morphometric and structural parameters related to cancellous and cortical bone architecture were measured (35).

### Transplantation

To purify LT-HSCs for transplant, enrichment of LT-HSCs from WT B6.SJL mice was performed using a biotinylated anti-”lineage” antibody cocktail (anti-CD3, CD4, CD8, CD11b, CD19, Gr1, NK1.1, and Ter119) followed by magnetic separation using EasySep Positive Selection kit (Stem Cell Technologies). After enrichment, cells were stained with streptavidin-Pacific Blue and antibodies to Sca1, cKit, CD150 and CD48. Cells were aseptically sorted on the BD FACS Aria II cell sorter to a 90.3-100% purity. Equivalent numbers of LT-HSCs (minimum of 225) were injected retro-orbitally into sublethally-irradiated (750 rads with a ^137^Cs source, J.L. Shepherd, Inc.) control B6 or *Sost*^*-/-*^ recipients under isoflurane anesthesia. Animals were supplemented with neomycin in the drinking water for 14 days post-transplant and analyzed for chimerism in the peripheral blood every 6 weeks, as described (33). Recipient mice were 16-22 weeks of age at the time of transplantation, and their bone marrow was analyzed at 16-20 weeks post-transplantation.

### Quantification of cytokines

Long bones were dissected and cleaned of muscle, and epiphyses removed. The bones were then place in a 0.5 mL tube in which a hole was introduced in the bottom using an 18 g needle.one marrow serum was harvested by removing epiphyses from both long bones after dissection and removal of muscle, placing bones in a 0.5 mL tube with 60 µL PBS with a hole in the bottom from 18 gauge needle. To harvest the BM serum, the 0.5mL tube containing the bone was then placed into a 1.5 mL microcentrifuge tube, and 60 µL PBS added to the bone. Tubes were centrifuged together for 30 seconds at 10,000 x g, collecting the serum supernatant in the 1.5 ml tube. Spleen serum was harvested by placing the spleen into a 1.5 mL tube with 60 µL PBS and homogenizing for 1 min. The sample was centrifuged for 30 sec at 10,000 x g, the serum supernatant collected. For peripheral blood collection, mice were heated gently under a heat lamp, and 7-8 drops of blood was collected from the tail veins in non-heparinized tubes. Blood was allowed to clot for at least 30 minutes and then was centrifuged for 10 minutes at 1,000 x g to separate the serum from the blood cells. Concentrations of cytokines (IL-1α, IL-1β, IL-6, IL-10, IL-12p70, IL-17A, IL-23, IL-27, MCP-1, IFN-β, IFN-γ, TNF-α, and GM-CSF) were determined from the serum of bone marrow, spleen and blood using the LEGENDplex™ Mouse Inflammation Panel from Biolegend, Inc. according to the manufacturer’s instructions.

### Hematoxylin and eosin (H&E)

Spleen samples were placed in 10% buffered formalin for 24 hours. Samples were then dehydrated with increasing concentrations of ethanol, cleared with xylene, then placed in wax and embedded in a mold. Samples were then sliced via microtomy, placed on slides and de-paraffinized before staining with hematoxylin followed by eosin.

### Complete blood counts

Peripheral blood was collected either by femoral vein blood collection immediately following cervical dislocation, or tail vein blood collection following heating under a heat lamp. In both cases, blood was collected in heparinized tubes. Complete blood counts were evaluated within 6 hours after collection on a Hemavet 950 Veterinary Hematology System.

## RESULTS

### Sclerostin-depleting antibody treatment changes hematopoietic differentiation

*Sost*^*-/-*^ mice display a drastic reduction in the BM cavity volume (36), which we hypothesized affects the hematopoietic stem cell niches within the BM (37) (38). Given the known relationship between Wnt signaling on hematopoietic stem cell (HSC) self-renewal and function (39), and that sclerostin is a Wnt signaling antagonist (3) (40), we hypothesized that removal of sclerostin through administration of neutralizing antibodies would promote HSC self-renewal. To test if acute depletion of SOST changed hematopoiesis, we performed studies in which SOST was depleted using sclerostin-specific antibodies (Scl-Ab), administered subcutaneously in 8-week-old mice in multiple doses for 6 weeks (Figure 1A). Scl-Ab treatment resulted in increased trabecular volume BV/TV and midshaft cortical thickness, similar to that observed in the *Sost*^-/-^ mice (Figure 1B, 1C and 1D) (41, 42). However, the BM total cellularity was unchanged (data not shown). Analysis of B cell development in the BM of Scl-Ab treated mice showed an increase in B cell progenitors (Hardy Fractions B-C and D), followed by a decrease in the mature recirculating B cells (Hardy Fraction F) (Supplemental Figure 1E), similar to the altered B cell development observed in the BM of 12-15 week old *Sost*^-/-^ mice (33). In the spleens of the Scl-Ab treated mice, a significant decrease in mature B cell populations (CD19^+^B220^+^ and IgM^+^B220^+^) was observed (Supplemental Figure 1F).

**Figure 1.**
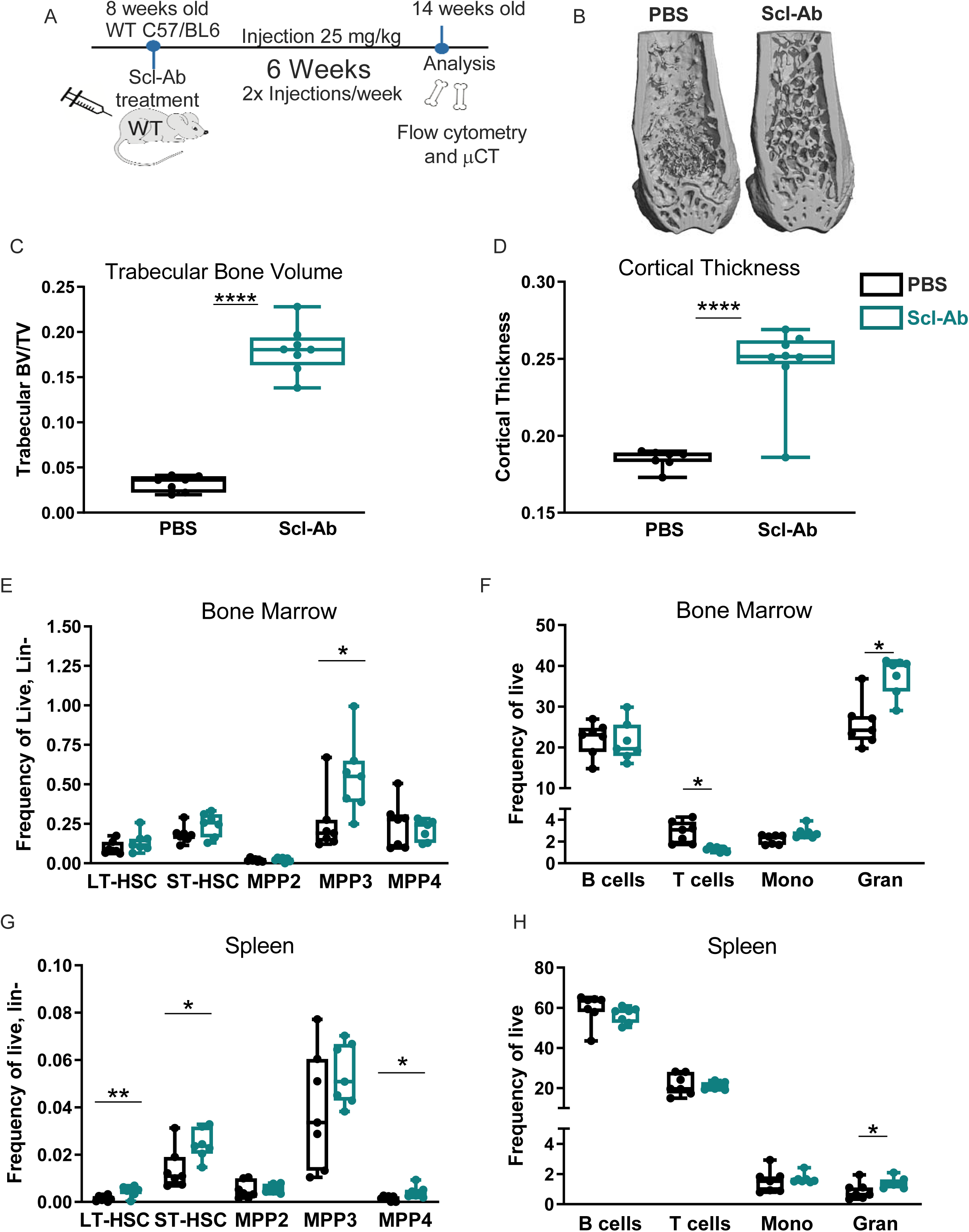
Sclerostin-depleting antibodies change hematopoietic differentiation. (A) Experimental scheme for Scl-Ab treatment and analysis; (B) Micro-CT images of femurs from PBS and Scl-Ab treated mice.; (C) Trabecular BV/TV; (D) Cortical bone mineral density; (E) Bone marrow frequency of LSK progenitors; (F) Bone marrow frequency of mature lineages; (G) Splenic frequency of LSK progenitors; (H) Splenic frequency of mature lineages. Female mice of 8 weeks of age were used for this study (n=7 for each group). p<0.05*, p<0.01**, p<0.001***,p<0.0001**** Mann-Whitney, non-parametric t-test.

The frequency of the HSPC subpopulations (LT-HSC, ST-HSC, MPP2, MPP3 and MPP4) was unaffected in the BM of Scl-Ab treated mice, with the exception of a slight increase in the frequency of MPP3 (a myeloid-biased progenitor) (Figure 1E). This change in frequency, however, did not result in a difference in MPP3 cell numbers (Supplemental Figure 1A). In the BM, CD3^+^ T lymphocytes were reduced and CD11b^+^ Gr1^+^ granulocytes were increased in both frequency and absolute number after Scl-Ab treatment (Figure 1F and Supplemental Figure 1B), consistent with the increase in MPP3 frequency. To test if these hematopoietic changes were confined to the bone, we also examined HSPC and hematopoietic lineages in the spleen. In contrast to the BM, the spleens displayed an increase in LT-HSCs, ST-HSCs and MPP4s (a lymphoid-biased progenitor) in frequency and cellularity (Figure 1G and Supplemental Figure 1C). The spleens also showed an increase in granulocytes in both cellularity and frequency (Figure 1H and Supplemental Figure 1D).

### Lack of sclerostin in the bone does not affect hematopoietic progenitor distributions in the bone marrow

Our observation that acute Scl-Ab treatment increased MPP3 distribution in the BM and LT-HSCs, ST-HSCs, and MPP4s in the spleen led us to inquire whether similar alterations would also be observed longer-term in a chronic *Sost* deficiency model. We elected to use a transplantation model as subtle changes in hematopoiesis in response to the bone microenvironment can be enhanced using transplantation approaches.

Equal numbers of purified wildtype (WT) CD45.1^+^ LT-HSCs (LSK, CD150^+^ CD48^-^) cells were transplanted into sub-lethally irradiated (750 rads) congenic CD45.2^+^ WT or *Sost*^-/-^ hosts (Figure 2A). The use of the congenic CD45.1 and CD45.2 mice is an established approach that permits the tracking of donor-derived (CD45.1^+^) hematopoiesis separate from host-derived (CD45.2^+^) hematopoiesis after transplant. This model also allows for comparison of fully developed LT-HSCs isolated from the same source of adult WT mice and their engraftment and differentiation after transfer into *Sost*^*-/-*^ or control microenvironments. Sixteen weeks after LT-HSC transplantation, the recipients were analyzed for BM and spleen cellularity, donor hematopoietic chimerism and the frequencies of donor-derived hematopoietic progenitors. As expected, total bone marrow cellularity was significantly decreased in the *Sost*^-/-^ recipients (Figure 2B (33)). Donor % hematopoietic chimerism within the BM was achieved at similar levels in WT and *Sost*^-/-^ recipients (Figure 2C), indicating no effect of the *Sost*^-/-^ microenvironment on donor WT LT-HSC engraftment after transplantation. However, the frequency of donor derived LSKs was increased in the BM (Figure 2D). We further analyzed the types of HSPCs in the chimeras for LT-HSCs, ST-HSCs, MPP2, MPP3, and MPP4 subpopulations (Figure 2E and 2G). Within the BM of WT→*Sost*^*-/-*^ chimeras, no statistically significant changes in proportions or absolute numbers of any HSPC subpopulations were observed (Figure 2E and Supplemental Figure 2A). Furthermore, no change in the expression of self-renewal and cell cycle genes *p21Cip1* and *HoxB4* were observed in *Sost*^*-/-*^ LT-HSCs sorted from the BM (Supplemental Figure 2C), and no evidence of changes in LSK quiescence or enhanced cell-cycling after 5-fluorouracil treatment was observed in *Sost*^*-/-*^ mice (Supplemental Figure 2D, 2E, 2F and 2G). In the BM of non-transplanted *Sost*^-/-^ mice, we observed a decrease in early-apoptotic LSKs, but no change in Ki67+ proliferating LSKs. (Figure 5C, 5E). However, similar to Scl-Ab treated mice, in LT-HSC and ST-HSC frequencies and absolute numbers in the spleen were increased in the WT→ *Sost*^*-/-*^ chimeras (Figure 2G and Supplemental Figure 2B). In addition, absolute numbers of MPP2 and MPP3 in the spleen were also increased (Supplemental Figure 2B).

**Figure 2.**
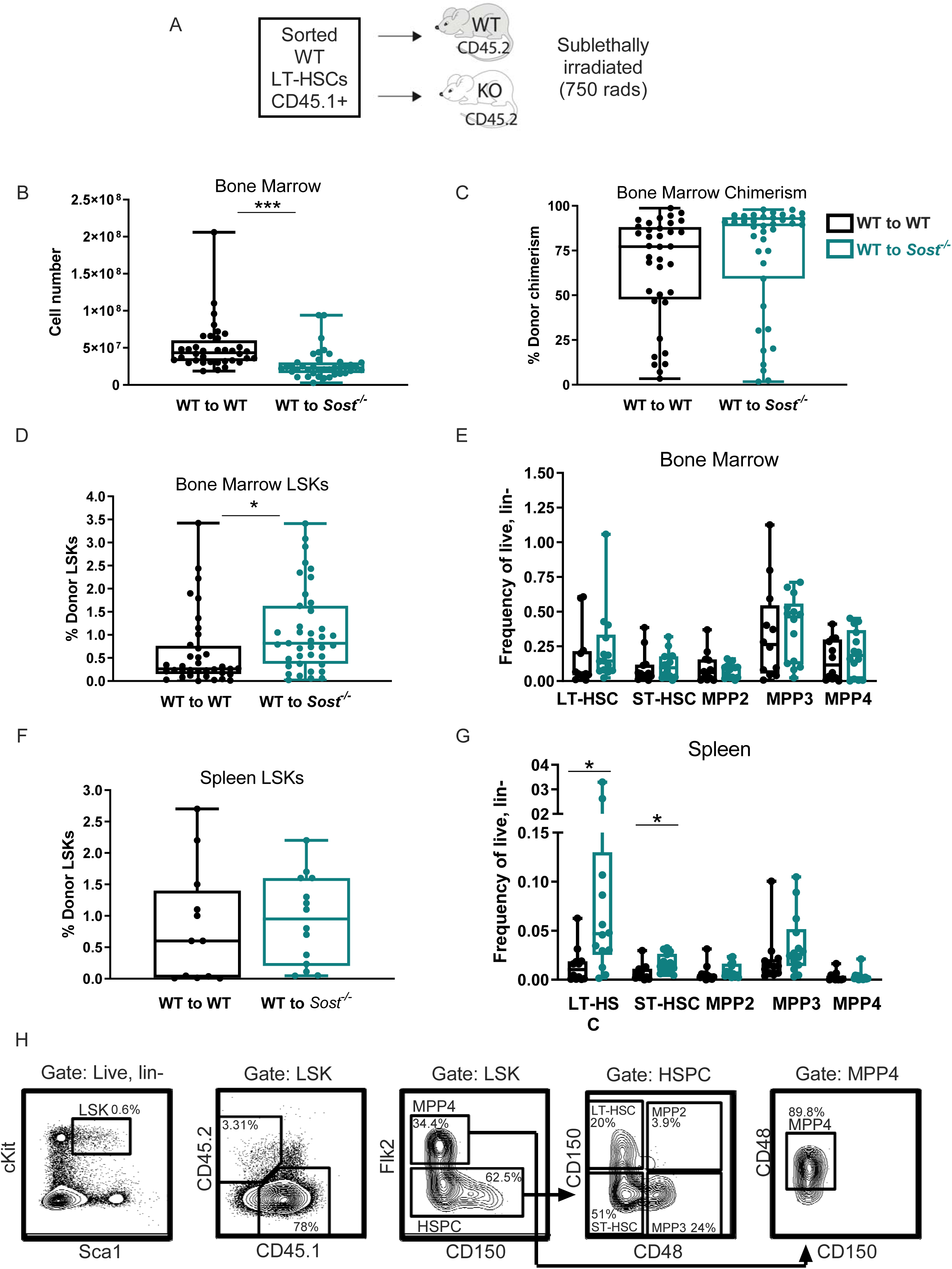
Lack of sclerostin in the bone does not alter hematopoietic progenitor distribution in the bone marrow. (A) Experimental scheme of LT-HSC transplant model to study hematopoiesis long-term in *Sost*-deficient BM microenvironments; (B) Total bone marrow cellularity in WT→WT (control) and WT→*Sost*^*-/-*^ chimeras; (C) Percent donor chimerism; (D) Frequency of donor-derived BM LSKs; (E) Frequency of donor derived BM HSPCs; (F) Frequency of donor derived splenic LSKs; (G) Frequency of donor derived spleen HSPCs; (H) Representative FACS plots depicting gating strategy for HSPC gating in chimeras. Ages of the BM donor mice ages ranged from 9-17 weeks. Ages of the recipients ranged from 16-22 weeks, when the phenotype of the *Sost*^*-/-*^ BM is already established. Both male and female donors and hosts were used. Donor LT-HSC cell numbers ranged from 235-400 cells per mouse. Sort purities ranged from 90.3% - 100%. Analysis of BM and spleens was performed 19-29 weeks post-transplantation. Data shown are compiled from 4 independent experiments. p<0.05*, p<0.01**, p<0.001***, p<0.0001**** Mann-Whitney, non-parametric t-test.

### Lack of sclerostin in the bone microenvironment results in a myeloid bias

We also analyzed hematopoietic differentiation broadly in the chimeras, using an antibody cocktail to quantify CD19^+^ B lymphocytes, CD3^+^ T lymphocytes, CD11b^+^ Gr1^-^ monocytes, and CD11b^+^ Gr1^+^ granulocytes (Figure 3A, 3B, 3C, 3D and 3E). In the BM, the frequency and number of donor-derived T cells and donor-derived B cell numbers were decreased in WT→*Sost*^*-/-*^ chimeras compared to WT→WT controls (Figure 3A and 3B). In addition, a significant increase in the frequency of donor-derived monocytes and granulocytes was observed in *Sost*^-/-^ hosts, but with similar absolute numbers as controls. (Figure 3A and 3B). Therefore, even though the MPP2, MPP3 and MPP4 HSPC progenitor frequencies were unchanged (Figure 2E), our observations of increased frequency of monocytes and granulocytes in the BM are consistent with skewing of LT-HSC hematopoietic differentiation towards the myeloid linages in the *Sost*^-/-^ BM microenvironment. Similarly, in the spleens of WT→ *Sost*^-/-^ chimeras, we observed an increase in the number of splenic MPP2 and MPP3 cells (Supplemental Figure 2B), accompanied with increased monocyte and granulocyte frequency and granulocyte cellularity in the WT→*Sost*^*-*/-^ spleens (Figure 3C, 3D).

**Figure 3.**
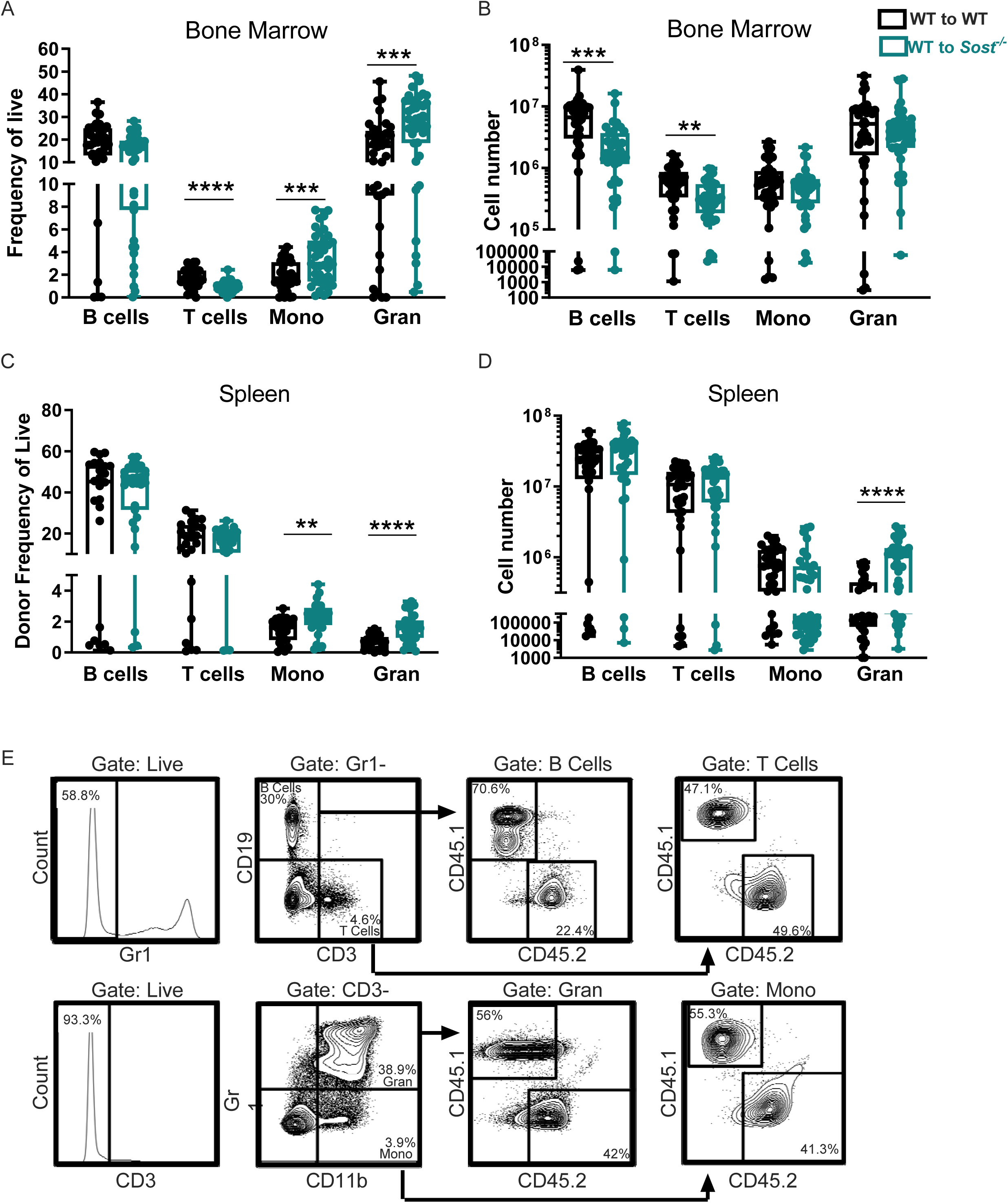
Lack of sclerostin in the bone microenvironment results in a myeloid bias. (A) Frequencies of donor-derived BM lineage populations in chimeras; (B) Cellularity of donor-derived BM lineage populations; (C) Donor-derived spleen lineage populations; (D) Cellularity of donor spleen lineage populations in chimeras; (E) Representative FACS plots depicting gating strategy for mature lineages in chimeras. Data shown is from a WT →*Sost*^*-/-*^ chimera. Data are from the same experiments described in Figure 3. p<0.05*, p<0.01**, p<0.001***, p<0.0001**** Mann-Whitney, non-parametric t-test.

### Sost^-/-^ mice express high levels of inflammatory cytokines in the bone marrow

CXCL12 is a chemokine that can influence HSC retention in the BM (43), and SCF is important for HSC maintenance. To further investigate the effects of *Sost* on the bone microenvironment, we analyzed the expression of CXCL12 and SCF in mesenchymal stem cells (MSCs), osteoblasts (OBs), endothelial cells (ECs), and other CD45-bone cells after collagenase-digestion of WT and *Sost*^-/-^ tibiae and femora. The distribution of MSCs, OBs and ECs differed in the Sost-/-mice compared to controls (Supplemental Figure 5D). We observed a lower expression of CXCL12 in the whole *Sost*^*-*/-^bone compared to controls, but this decreased expression could not be attributed to MSCs, OBs or ECs (Supplemental Figure 5B). In addition, SCF levels were similar in *Sost*^*-*/-^ and control bones (Supplemental Figure 5C).

The immune system is very responsive to inflammatory signaling caused by insults and environmental disturbances (44). Pietras *et al*. (45) and others have shown that inflammatory responses in the bone marrow are accompanied by changes in the blood system, including overproduction of myeloid cells and decrease in production of lymphoid cells. We hypothesized that the *Sost*^-/-^ BM microenvironments may express increased levels of pro-inflammatory cytokines. We assessed 13 different inflammatory cytokines in the serum of the *Sost*^-/-^ BM and spleen, and found TNF-α, MCP-1 and IL-1α to be significantly increased in the BM (Figure 4A). This increase was localized to the *Sost*^*-/-*^ BM as no difference in TNF-α, MCP-1 and IL-1α was observed in the spleen (Figure 4B) or peripheral blood (data not shown). No difference in IL-1β, IL-6, IL-10, IL-12p70, IL-17A, IL-23, IL-27, IFN-β, IFN-γ, and GM-CSF were observed in any tissue examined (data not shown).

**Figure 4.**
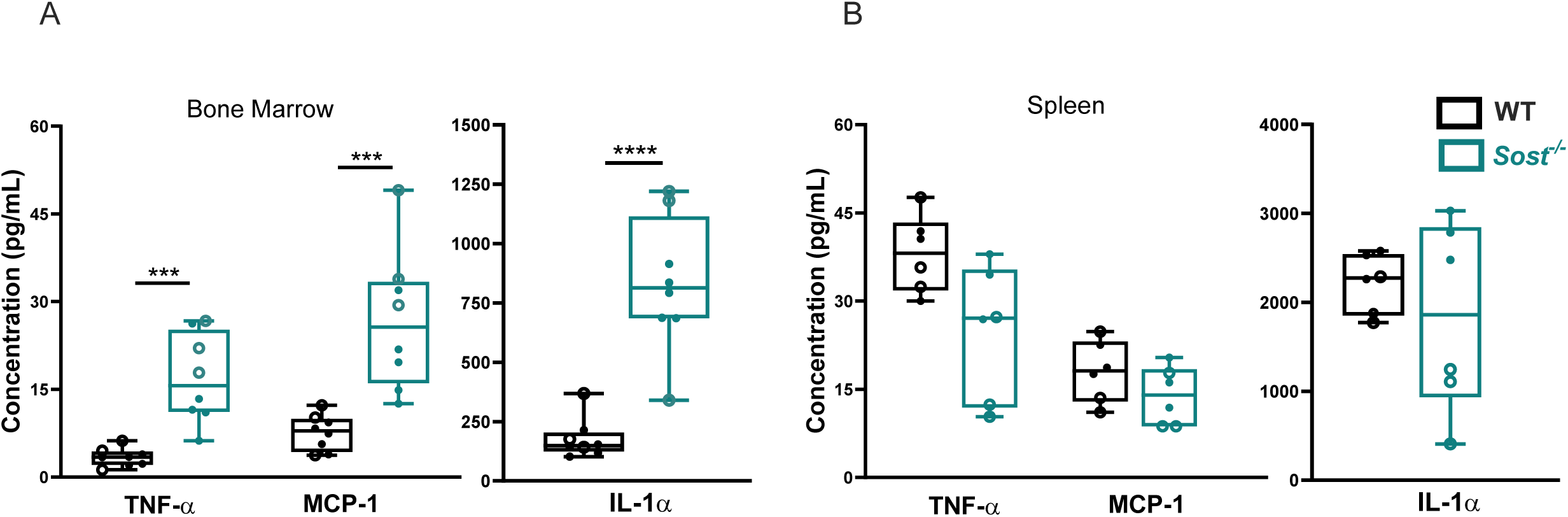
Elevated levels of inflammatory cytokines in Sost^-/-^ bone marrow. (A) TNFα, MCP-1 and IL-1α concentrations in WT and *Sost*^*-/-*^ bone marrow; (B) Analysis of inflammatory cytokine concentrations in WT and *Sost*^*-/-*^ spleen. Open circles represent male mice and filled circles represent female mice. Male (4 WT and 4 *Sost*^*-/*-^) and female (4 WT and 4 *Sost*^*-/*-^) mice ranging in ages 25-36 weeks were used. All samples were analyzed using a single LEGENDPLEX kit. p<0.05*, p<0.01**, p<0.001***, p<0.0001**** Mann-Whitney, non-parametric t-test.

### Evidence of extramedullary hematopoiesis in Sost^-/-^ mice

Insults to the BM environment, such as inflammation (46-49), can result in extramedullary hematopoiesis (EMH) in the spleen. The increase in the splenic HSPC compartment indicated EMH was prominent in the spleens of WT→*Sost*^*-/-*^ chimeras. Consistent with this, a significant increase in the frequency and cellularity of splenic LT-HSC and ST-HSCs was evident (Figure 2G and Supplemental Figure 2B). Splenic MPP2 and MPP3 populations were also increased in cellularity (Supplemental Figure 2B). To determine the kinetics of the onset of EMH, we analyzed the LT-HSC frequency in the spleens of unmanipulated *Sost*^*-/-*^ mice as a function of age, and observed that LT-HSCs in *Sost*^*-/-*^ spleens were significantly increased by 34 weeks of age, whereas the increase in ST-HSCs began as early as 15 weeks (Supplemental Figure 3D, 3E, 3F, 3G). The spleen contains an increase in proliferating LSK HSPCs in active cell cycle, as measured by Ki67 expression (Figure 5B and 5F). Paradoxically, an increase in early-apoptotic LSKs was also observed in *Sost*^-/-^ spleens. The size of the spleens of *Sost*^*-/-*^ mice was visibly enlarged compared to controls (Supplemental Figure 3A, 3C), and we observed a trend toward an increase in the splenic red pulp area, where LT-HSCs and red blood cells are found in EMH (46) (Supplemental Figure 3B, 3C). To test their hematopoietic function, we purified splenic HSPCs from *Sost*^*-/-*^ mice and transplanted them into sublethally irradiated WT hosts (Supplemental Figure 4A). It is important to note there was no difference in hematopoietic engraftment or differentiation amongst HSPCs isolated from WT BM, *Sost*^*-/-*^ BM or *Sost*^*-/-*^ spleens (Supplemental Figure 4D-4G). Taken together, these data indicate EMH is elevated in the *Sost*^*-/-*^ mice but that the microenvironment of the *Sost*^*-/-*^ spleen does not seem to permanently harm the engraftment or differentiation capability of HSPCs.

**Figure 5.**
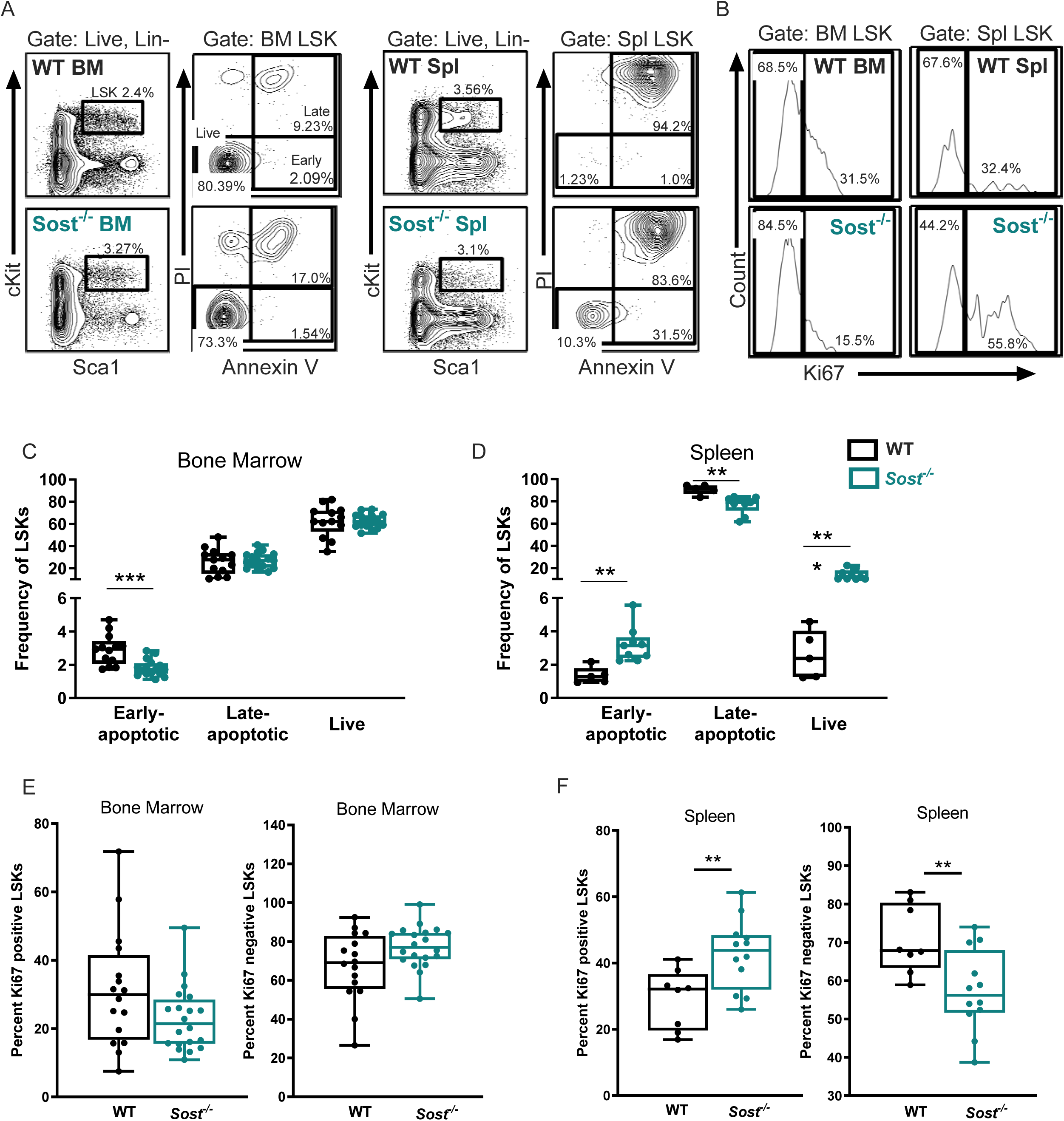
BM and splenic LSKs in Sost-/-mice display opposite patterns of proliferation and early apoptosis. (A) Representative FACS analysis of LSKs stained with Annexin V and PI, showing live cells, and cells in early and late apoptosis; (B) FACS analysis of LSKs stained with Ki67; (C) Summary of apoptosis staining results in the LSKs of the BM; (D) Summary of apoptosis staining results in the LSKs of the spleen; (E) Frequencies of Ki67+ BM LSKs; (F) Frequencies of Ki67+ splenic LSKs. Data on apoptosis was compiled from 2 independent experiments, using male mice with ages ranging from 6-63 weeks with age matched controls in the same experiment. Data on proliferation was compiled from 3 independent experiments, using male and female mice with ages ranging from 23-54 weeks. p<0.05*, p<0.01**, p<0.001***, p<0.0001**** Mann-Whitney, non-parametric t-test.

### Erythrocyte development is altered in Sost^-/-^ mice

Chronic inflammation has been shown to induce anemia (50), and we hypothesized the increase in TNF-α, MCP-1 and IL-1α would result in a decrease of mature erythrocytes (red blood cells, RBC) in the BM of *Sost*^-/-^ mice. To determine if mature RBCs and erythrocyte progenitors in the BM were reduced, we performed flow cytometry analysis with Ter119 and CD71 cellular markers, which allows for discrimination of developmentally distinct RBC progenitor populations (Stage 1: CD71^+^Ter119^-^, Stage 2: CD71^+^Ter119^+^, Stage 3: CD71^mid^Ter119^+^, and Stage 4: CD71^-^ Ter119^+^) (51). The most immature progenitors (Stage 1) express intermediate levels of Ter119 and high levels of CD71 and as they mature, begin to downregulate CD71 and upregulate Ter119 (Figure 6A). In *Sost*^-/-^ BM we observed a primary increase in Stage 2 RBC frequency with an immediate decrease in Stage 3 (Figure 6B). Interestingly, in the spleen, we observed that Stages 1-3 are increased, and Stage 4 is decreased, indicating a possible RBC developmental block (Figure 6C).

**Figure 6.**
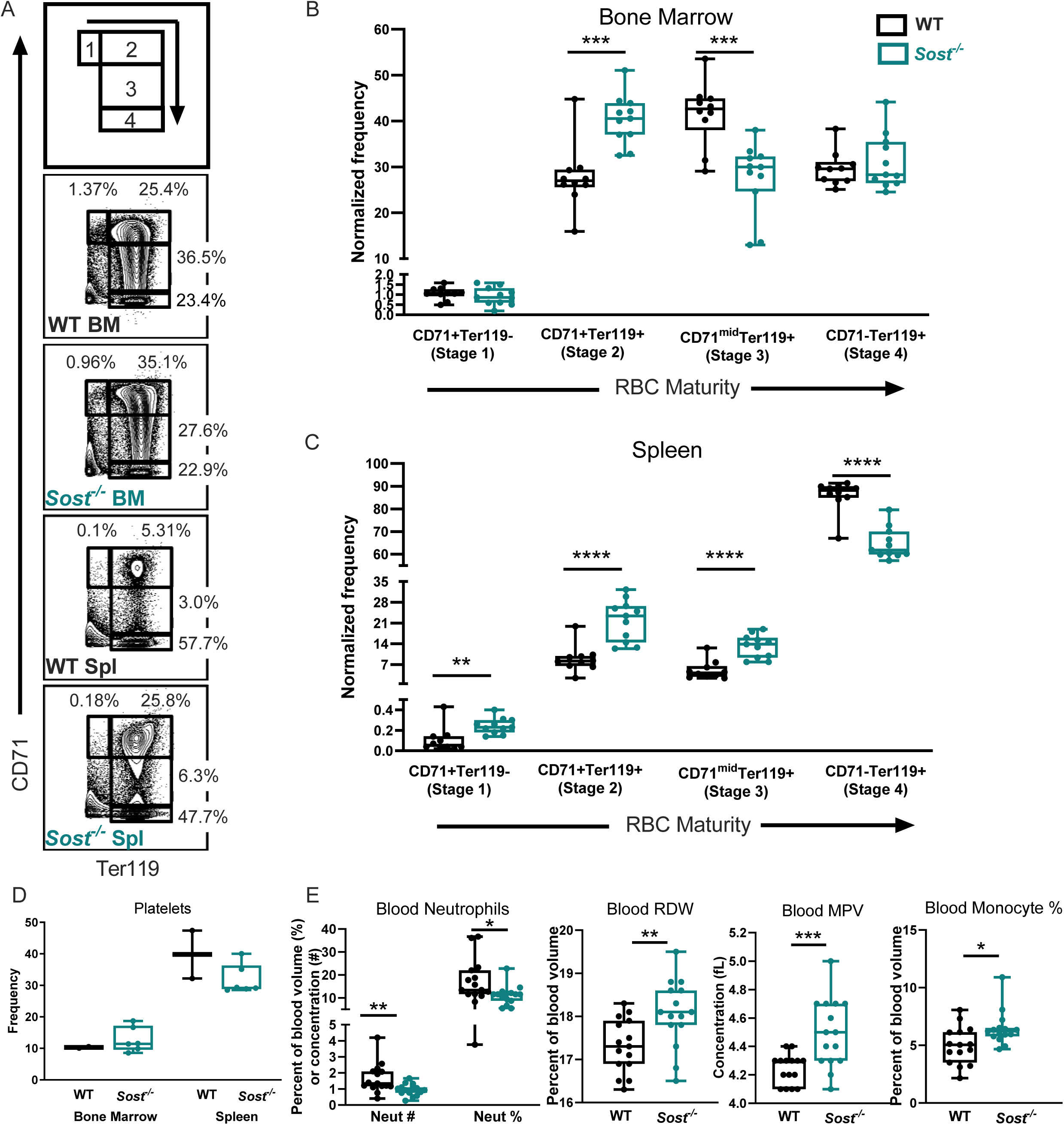
RBC development in the Sost^-/-^ BM is altered and developmentally blocked in the Sost^-/-^ spleen. (A) Representative FACS plots depicting stages of RBC development (B) Normalized frequencies of Stages 1-4 bone marrow RBC progenitor populations (C) Normalized frequencies of Stages 1-4 splenic RBC populations (D) Bone marrow platelet frequency measured by flow cytometry (E) CBC analysis depicting neutrophil concentration and cellularity, red blood cell distribution width (RDW), mean platelet volume (MPV). Data on RBC development in the BM and spleen mice were compiled from 3 independent experiments, and male and female mice of 9-60 weeks of age were used. Platelet analysis in the BM and spleen were compiled from 2 independent experiments, using mice of both sexes ranging in age from 6-60 weeks. Peripheral blood data were compiled from 2 independent experiments, using mice of both sexes ranging in age from 25-48 weeks. p<0.05*, p<0.01**, p<0.001***, p<0.0001**** Mann-Whitney, non-parametric t-test.

We utilized a Hemavet for complete blood cell analysis to further examine RBC differences in the *Sost*^-/-^ mice. This analysis revealed an increase in red blood cell distribution width (RDW) in *Sost*^-/-^ mice (Figure 6E). A rise in RDW values indicate greater variation in RBC size and shape and is known as anisocytosis, which can be caused by anemia (52). We also examined megakaryocytes, which give rise to both RBCs and platelets and observed no differences in platelet frequency in the BM or spleens of *Sost*^-/-^ mice by flow cytometry (Figure 6D). However, we observed an increase in mean platelet volume (MPV) the peripheral blood of in *Sost*^*-/-*^ mice (Figure 6E). An increase in MPV is usually observed when there is destruction of platelets, commonly seen in myeloproliferative diseases. Interestingly, despite the increase in granulocytes in both the BM and spleen in Scl-Ab treated mice and WT→*Sost*^*-/-*^ chimeras, complete blood cell analysis of peripheral blood revealed a significant decrease in neutrophils (the major type of granulocyte) number and frequency, indicating a decrease in mobility of neutrophils out of the BM and spleen (Figure 6E). A comprehensive list of the CBC data is shown in Supplemental Table 2.

## DISCUSSION

The sclerostin monoclonal antibody romosozumab is now approved for treatment of osteoporosis in the US, Japan, Canada, Australia, and S. Korea) (53, 54). This treatment is effective at treating osteoporosis; however, our previous studies and our current study in Scl-Ab treated mice and studies in hematopoietic differentiation in older *Sost*^*-/-*^ mice and long-term studies in WT→*Sost*^*-*/-^ chimeras suggest that this treatment may have unintended effects on immune development of patients, many of whom are older and have less plastic immune systems (55).

In our models of acute and chronic *Sost* depletion, we observed alterations in hematopoietic lineage differentiation, favoring myelopoiesis and decreasing lymphopoiesis in the BM. Granulocytes were markedly increased in both models, but monocytes were increased only in the chronically depleted, WT→*Sost*^-/-^chimeras. One possible explanation for this is that monocyte changes may take longer to emerge and may not be observable in an acute model of sclerostin depletion. Our previous work did not indicate any changes in the frequencies of hematopoietic progenitors within the Lin-Sca1^high^ c-Kit^high^ (LSK) compartment in WT→*Sost*^-/-^chimeras (33), but in our earlier work, the chimeras were analyzed within a shorter, five-week time-course, which is not sufficient to assess the behavior of LT-HSCs. In any case, the similar phenotypes of increased myelopoiesis and decreased lymphopoiesis in the SOST-depleted or *Sost*-deficient BM strongly suggests that blocking SOST could contribute to “inflammaging”, defined as chronic, sterile, low-grade inflammation that contributes to the pathogenesis of age related diseases (56). Increased inflammation during aging can result in concurrent decline in HSC function. The decline of HSC functionality includes a reduction in long-term repopulating potential, homing and engraftment after transplant (19), a decrease in lymphopoiesis and increase in myelopoiesis (20, 21) as well as an increase in total number of LT-HSCs (22), similar to what we observed *Sost*^-/-^ mice. Chronic inflammation has also been associated with aging, characterized by anemia, immunosenescence, thrombocytosis (23) as well as overproduction of inflammatory cytokines IL-1, TNF-α and IL-6 (24). Consistent with this, *Sost*^-/-^ mice display significantly higher levels of TNF-α, and IL-1a, and we also observed high levels of MCP-1. The high levels of MCP-1 may explain the increase in monocytes observed in the *Sost*^-/-^ mice, as MCP-1and its receptor CCR2 play critical roles in monocyte recruitment during inflammation and bone remodeling (57, 58). It has also been suggested that MCP-1 plays a critical role in the recruitment of monocytes to the bone, as it is induced during osseous inflammation (59).

Another feature of inflammaging is in the induction of EMH in the spleen, where we also observed an increase in myelopoiesis. The decreased expression of CXCL12 in the BM of *Sost*^*-/-*^ mice suggests that LT-HSCs may be unable to be retained in the BM and are actively migrating to the spleen. Altogether, our data indicate that SOST depletion and *Sost*-deficiency not only affects B cell development in the BM relatively early, but also affects monocyte and granulocyte differentiation and creates an inflammatory microenvironment, that may become more severe over time.

We sought to determine if the changes in lineage development downstream of the LT-HSCs could be explained by changes in their proliferation and apoptosis. It was interesting to observe an increase in early apoptosis in *Sost*^*-/-*^ BM LSKs, indicating that maintenance of cell membrane integrity was intact, with no change LSKs in late apoptosis/necrosis, which lack intact cell membranes. Apoptosis is an active, energy dependent process with apoptotic cells expressing surface markers that permit their ingestion by phagocytosis (60). Due to the deterioration of the cell membrane after late apoptosis, most cells are cleared by phagocytes with their cell membranes still intact before they release their potentially inflammatory intracellular contents. The increase in monocytes in the Sost^-/-^ BM may result in an increase in phagocytosis of early apoptotic cells where the cell membrane is still intact. The observation of increased frequency of proliferative, live LSKs and decrease in LSKs in late apoptosis stages in the spleen is consistent with the EMH observed in the *Sost*^-/-^ mice. Paradoxically, the splenic LSKs also displayed an increase in early apoptosis. Further investigation is required to reconcile this.

Our analysis of erythrocyte development in the *Sost*^*-*/-^ mice revealed a significant increase in RDW, known as anisocytosis (61). RDW is elevated in folate deficiencies characteristic of macrocytic anemia (62). Although folate levels in *Sost*^-/-^ mice remains untested, decreased folate levels have been associated with poor bone health, in humans (63). Anisocytosis can be split into two groups, anisocytosis with microcytosis, characterized by low iron (e.g. sickle cell anemia), and anisocytosis with macrocytosis, characterized by folate deficiency (e.g. myelodysplastic syndrome). We know that *Sost*^-/-^ mice have normal hemoglobin levels (Supplemental Table 2) indicating no issues with iron; however, iron levels in *Sost*^-/-^ mice have not been directly quantified to date. The increase in mean platelet volume (MPV) in the blood is usually associated with an increase in platelet production, or destruction of platelets as seen in myeloproliferative diseases (64-66) such as chronic myeloid leukemia (CML), myelofibrosis, and myelodysplastic syndrome (MDS) (67-69). *Sost*^-/-^ mice displayed no changes in the blood platelet levels (data not shown). Future studies are needed to investigate if increased MPV is related to the increased myelopoiesis observed in *Sost*^*-/-*^ mice.

The significant decrease in mature splenic RBCs in *Sost*^-/-^ mice may represent a direct consequence of inflammaging. Prior studies have shown that TNF-α inhibits erythropoiesis directly through activation of GATA-2, whose over-expression is known to inhibit erythropoiesis in favor of megakaryopoiesis (70) (71). Reduced capacity for erythropoiesis or platelet production is often compensated by extramedullary hematopoiesis in the spleen (72). Indeed, we observed a significant increase in immature RBC production in the spleen of the *Sost*^-/-^ mice. The induction of EMH in the spleens of *Sost*^-/-^ mice with age seems to suggest an attempt at compensation for the decrease in RBC production in the spleen. As *Sost* is not expressed in the spleen, the physiological mechanisms that drive extramedullary hematopoiesis in the spleens of the *Sost*^-/-^ mice is of particular interest to our laboratory. Certainly, the biology of splenic hematopoietic niches is not well understood, and future studies could determine if LT-HSCs in *Sost*^*-/-*^ mice are actively migrating from the BM and into the spleen or if LT-HSCs are expanding in the spleen de novo. Nevertheless, we determined no differences in *Sost*^*-/-*^ splenic and BM LT-HSC function after transplantation. In addition, future studies to understand the identify and test the function of stromal cells in the spleen that drive extramedullary hematopoiesis in *Sost*^-/-^ mice.

In summary, our studies extend previous studies and reveal novel information on the effects of sclerostin-deficiency in bone and sclerostin-depleting treatments on hematopoietic stem cells. This information may be useful in monitoring humans treated with romosozumab for changes in immune cell frequencies, chronic inflammation, and signs of anemia, such that treatments for osteoporosis can be modified to address these hematopoietic changes.

## ACKNOWLEDGMENTS

This work was supported by University of California, Merced faculty research funding, the UC Cancer Research Coordinating Committee grant CRR-13-201415, and NIH grant R15HL121786 (J.O.M.) and student research grants to AM and MC. GL works under the auspices of the U.S. Department of Energy by Lawrence Livermore National Laboratory under Contract DE-AC52-07NA27344. Authors’ roles: AE and GL provided the Sost^-/-^ mice to begin our colony at UC Merced. Study design: CD, GL and JOM. Study conduct: CD and JOM. Data collection and data analysis: CD, BC, MC, BF, SE, AM, AR, DM, JOM. Drafting manuscript: CD and JOM. Revising manuscript content: CD, GL, JOM. Approving final version of manuscript: JOM. JOM takes responsibility for the integrity of the data analysis. The authors thank Dr. Aris Economides of Regeneron, Inc. for providing Scl-Ab, and the staff of the Department of Animal Research Services for excellent animal care, and Dr. David Gravano of the Stem Cell Instrumentation Foundry at UC Merced for outstanding technical support in flow cytometry and histology.

