## Supplemental Figures for "Sclerostin depletion induces inflammation in the bone marrow of mice"

**SUPPLEMENTARY MATERIAL**

**Supplemental Materials and Methods**

*Bone digestion for quantitative real-time PCR.* Bone digests to obtain MSC, OBs, ECs and enriched osteocytes was performed similar to Schepers et al. (1). Femurs and tibias were harvested in 1x HBSS without serum and crushed using a mortar and pestle. Bone marrow was washed away and the bone chips were then incubated with 3mg/ml Type I Collagenase in HBSS without serum (Worthington) at 37°C on a shaker at 110 rpm for 1 hour. After incubation the supernatant was harvested from the bone chips, filtered through a 70 um cell strainer, and then washed with 1x HBSS-2% FCS to stop the action of the collagenase. Cells were centrifuged at 300-500 x g for 5 minutes to wash, resuspended in 1x HBSS-2% FCS, and cell counts performed using Trypan Blue staining on a hematocytometer. Cells were then stained for flow cytometry with antibodies to lineage-positive cells (CD3, CD4, CD8, CD11b, CD19, NK1.1, GR1 and TER119, all in PE-Cy7), CD45-eFluor450, Sca1-FITC, CD31-APC, CD51-biotin followed by streptavidin-PE. PI was added as viability marker. Cells were sorted using a FACS Aria III and cell pellets stored for real-time PCR.

*Real-time PCR.* LT-HSCs were purified from pooled BM from control and *Sost*$^{-/-}$ long bones by flow cytometry, as described in the main text. Cells were pelleted and resuspended in RNeasy Lysis Buffer with 2-mercaptoethanol (Qiagen). Total RNA was purified using Qiagen RNeasy Micro Kit (Qiagen) according to manufacturer's protocol. RNA concentration and purity was analyzed using a NanoDrop ND-1000 Spectrophotometer (Thermo Fisher Scientific). The same amount of RNA from each sample was mixed with qScript XLT-1 Step, RT-qPCR ToughMix (Quantabio Cat. 66149433) together with specific TaqMan expression assays. Real-time quantitative PCR was ran on an Applied Biosystems thermocycler in this sequence: 1 cycles at 50°C for 10 minutes for cDNA synthesis, 1 cycle at 95°C for 1 minute for initial denaturation and then 40 cycles of amplification at 95°C for 5 seconds then 60°C for 45 seconds, using QuantStudio 3 software (ThermoFisher). The following TaqMan gene expression assays (Thermo Fisher Scientific) were used: HoxB4-FAM (Mm00657964_m1), p21/Cip1/Cdkn1a-FAM (Mm00432448_m1), and GAPDH-VIC (MM999915_g1). Expression of HoxB4 and p21/Cip1 in KO LT-HSCs relative to the average WT gene expression was calculated using the double delta Ct method (2). RNA from digested bones and qPCR for CXCL12 and SCF was performed as described (3).

*Cell cycle analysis after 5-fluorouracil treatment.* B6 and *Sost*$^{-/-}$ mice (16-18 weeks of age) were injected with PBS or 100mg/kg 5-fluorouracil (Sigma) intraperitoneally. Seven days later, mice were euthanized and bone marrow cells were harvested for flow cytometry as described above. First, cells were extracellularly stained with anti-Lineage, Sca-1 and cKit antibodies, and then fixed and permeabilized using the eBioscience™ Foxp3 / Transcription Factor Staining Buffer Set for intranuclear staining. Cells were stained on ice for 30 minutes in the dark with anti-Ki67-PE (clone SolA15) or isotype-PE control mAb, purchased from eBioscience. After incubation, cells were washed twice with permeabilization buffer, resuspended in 400µl FACS buffer and transferred to tubes for acquisition on the LSR II. DAPI was added to the samples at a final concentration of 1.0 ug/ml immediately before acquisition.

**Supplementary Table 1. Antibodies used for flow cytometry**

| **Specificity** | **Clone** | **Fluorochrome** |
|---|---|---|
| Annexin V | | APC |
| CD11b | M1/70 | Biotin |
| CD127 (Il7rα) | A7R34 | Biotin |
| CD127 (Il7rα) | A7R34 | PerCP Cy5.5 |
| CD150 | mShad | FITC |
| CD16/32 | 93 | PerCP-Cy5.5 |
| CD19 | 6D5 | Biotin |
| CD19 | 6D5 | PE |
| CD3 | 2C11 | Biotin |
| CD3 | 2C11 | APC |
| CD31 | 390 | APC |
| CD34 | RAM34 | FITC |
| CD4 | Gk1.5 | Biotin |
| CD41 | MWReg30 | BUV395 |
| CD45 | 30-F11 | APC-Cy7 |
| CD45 | 30-F11 | PerCP Cy5.5 |
| CD45.1 | A20 | FITC |
| CD45.1 | A20 | PE |
| CD45.1 | A20 | PerCP-Cy5.5 |
| CD45.2 | 104 | APC-Cy7 |
| CD45R (B220) | RA3-6B2 | Biotin |
| CD48 | HM48 | PE-Cy7 |
| CD48 | HM48 | APC |
| CD5 | 53-7.3 | Biotin |
| CD51 | RMV-7 | Biotin |
| CD8 | 53.6.7 | Biotin |
| cKit | 2B8 | APC |
| cKit | 2B8 | APC |
| F4/80 | BM8 | Biotin |
| Flk2 | A2F10 | PE |
| Gr1 | RB6-8C3 | Biotin |
| Gr1 | RB6-8C3 | PE-Cy7 |
| Ki67 | SolA15 | APC |
| NK1.1 | PK136 | Biotin |
| Sca1 | D7 | BV510 |
| Sca1 | D7 | FITC |
| Streptavidin | | PerCP Cy5.5 |
| Streptavidin | | PacBlue |
| Streptavidin | | PE |
| Ter119 | TER119 | Biotin |

### Supplementary Table 2. Complete blood count values

| Genotype & Sex | | WBC K/uL | Ne# K/uL | Ly# K/uL | Mo# K/uL | Eo# K/uL | Ba# K/uL | Ne% % | Ly% % | Mo% % | Eo% % | Ba% % | RBC M/uL | HB g/dL | HCT % | MCV fL | MCH Pg | MCHC g/dL | RDW % | PLT K/uL | MPV fL |
|---|---|---|---|---|---|---|---|---|---|---|---|---|---|---|---|---|---|---|---|---|---|
| | *control* | *17.32* | *9.83* | *6.15* | *0.39* | *0.16* | *0.79* | *56.73* | *36* | *2.26* | *0.95* | *4.55* | *9.09* | *16.3* | *52.6* | *57.9* | *17.9* | *31* | *26.1* | *967* | *7.6* |
| **WT Female** | 1 | 7.16 | 1.34 | 5.54 | 0.22 | 0.04 | 0.01 | 18.72 | 77.4 | 3.12 | 0.57 | 0.16 | 9.97 | 13.8 | 53.3 | 53.5 | 13.8 | 25.9 | 16.5 | 802 | 4.2 |
| | 2 | 9.02 | 1.21 | 7.12 | 0.58 | 0.09 | 0.02 | 13.44 | 78.9 | 6.38 | 1.04 | 0.22 | 10.18 | 14.4 | 53.1 | 52.2 | 14.1 | 27.1 | 17.4 | 739 | 4.4 |
| | 3 | 9.16 | 0.76 | 7.64 | 0.74 | 0.02 | 0.01 | 8.29 | 83.4 | 8.07 | 0.19 | 0.07 | 10.5 | 14.8 | 56.5 | 53.8 | 14.1 | 26.2 | 16.5 | 695 | 4.3 |
| | 4 | 10 | 1.17 | 8.27 | 0.51 | 0.03 | 0.01 | 11.73 | 82.7 | 5.09 | 0.32 | 0.13 | 10.8 | 14.7 | 55.6 | 51.5 | 13.6 | 26.4 | 17.2 | 614 | 4.4 |
| | 5 | 10.14 | 1.1 | 8.46 | 0.45 | 0.1 | 0.03 | 10.86 | 83.4 | 4.42 | 0.99 | 0.29 | 9.78 | 14.5 | 51.9 | 53.1 | 14.8 | 27.9 | 17.1 | 642 | 4.3 |
| | **Mean** | **9.096** | **1.12** | **7.406** | **0.5** | **0.06** | **0.02** | **12.61** | **81.2** | **5.42** | **0.62** | **0.17** | **10.25** | **14.4** | **54.1** | **52.8** | **14.1** | **26.7** | **16.9** | **698** | **4.32** |
| | **SD** | **1.19** | **0.22** | **1.17** | **0.19** | **0.04** | **0.01** | **3.89** | **2.81** | **1.89** | **0.38** | **0.08** | **0.409** | **0.39** | **1.9** | **0.95** | **0.45** | **0.803** | **0.42** | **75.3** | **0.08** |
| ***Sost*$^{-/-}$ Female** | 11 | 7.28 | 1.66 | 5.09 | 0.46 | 0.06 | 0.01 | 22.76 | 69.9 | 6.34 | 0.81 | 0.17 | 9.44 | 13.4 | 53.1 | 56.2 | 14.2 | 25.2 | 16.5 | 659 | 4.7 |
| | 12 | 5 | 0.27 | 4.34 | 0.32 | 0.04 | 0.02 | 5.47 | 86.9 | 6.35 | 0.79 | 0.5 | 8.72 | 13.3 | 47.4 | 54.4 | 15.3 | 28.1 | 18 | 549 | 4.7 |
| | 13 | 7.68 | 0.93 | 6.27 | 0.38 | 0.07 | 0.02 | 12.15 | 81.7 | 4.92 | 0.94 | 0.3 | 9.21 | 13.9 | 50.5 | 54.8 | 15.1 | 27.5 | 17.8 | 607 | 4.7 |
| | 14 | 4.26 | 0.49 | 3.26 | 0.46 | 0.03 | 0.01 | 11.6 | 76.5 | 10.9 | 0.82 | 0.23 | 8.77 | 14.2 | 47.8 | 54.5 | 16.2 | 29.7 | 18.8 | 606 | 5 |
| | 15 | 6.42 | 0.77 | 5.03 | 0.52 | 0.08 | 0.02 | 12.02 | 78.3 | 8.09 | 1.3 | 0.3 | 10.25 | 14.1 | 58.2 | 56.8 | 13.8 | 24.2 | 16.8 | 570 | 4.6 |
| | **Mean** | **6.128** | **0.82** | **4.798** | **0.43** | **0.06** | **0.02** | **12.8** | **78.6** | **7.32** | **0.93** | **0.3** | **9.278** | **13.8** | **51.4** | **55.3** | **14.9** | **26.94** | **17.6** | **598** | **4.74** |
| | **SD** | **1.465** | **0.53** | **1.105** | **0.08** | **0.02** | **0.01** | **6.233** | **6.29** | **2.3** | **0.21** | **0.12** | **0.622** | **0.41** | **4.44** | **1.09** | **0.95** | **2.226** | **0.93** | **42** | **0.15** |
| **WT Male** | 6 | 18.04 | 4.2 | 13.33 | 0.39 | 0.08 | 0.04 | 23.29 | 73.9 | 2.16 | 0.45 | 0.2 | 10.27 | 14.2 | 52.8 | 51.4 | 13.8 | 26.9 | 18.1 | 927 | 4.3 |
| | 7 | 16.38 | 2.57 | 13.15 | 0.57 | 0.07 | 0.01 | 15.72 | 80.3 | 3.5 | 0.42 | 0.09 | 10.14 | 13.7 | 52.5 | 51.8 | 13.5 | 26.1 | 17.5 | 887 | 4.3 |
| | 8 | 16.86 | 2.27 | 13.72 | 0.74 | 0.1 | 0.03 | 13.44 | 81.4 | 4.4 | 0.6 | 0.15 | 10.12 | 13.7 | 51.5 | 50.9 | 13.5 | 26.6 | 16.9 | 817 | 4.2 |
| | 9 | 17.98 | 2.09 | 14.9 | 0.91 | 0.07 | 0.01 | 11.61 | 82.9 | 5.05 | 0.4 | 0.07 | 10.86 | 14.9 | 57.2 | 52.7 | 13.7 | 26 | 17.3 | 746 | 4.1 |
| | 10 | 11.02 | 0.41 | 10.04 | 0.51 | 0.04 | 0.02 | 3.76 | 91.1 | 4.61 | 0.34 | 0.17 | 5.65 | 7.6 | 29 | 51.3 | 13.5 | 26.2 | 16.3 | 503 | 4.2 |
| | **Mean** | **16.06** | **2.31** | **13.03** | **0.62** | **0.07** | **0.02** | **13.56** | **81.9** | **3.94** | **0.44** | **0.14** | **9.408** | **12.8** | **48.6** | **51.6** | **13.6** | **26.36** | **17.2** | **776** | **4.22** |
| | **SD** | **2.905** | **1.35** | **1.804** | **0.2** | **0.02** | **0.01** | **7.057** | **6.18** | **1.15** | **0.1** | **0.05** | **2.122** | **2.96** | **11.2** | **0.68** | **0.14** | **0.378** | **0.67** | **168** | **0.08** |
| ***Sost*$^{-/-}$ Male** | 16 | 11 | 1.06 | 9.2 | 0.65 | 0.07 | 0.02 | 9.61 | 83.6 | 5.93 | 0.65 | 0.2 | 9.47 | 13.2 | 51.7 | 54.6 | 13.9 | 25.5 | 17.8 | 774 | 4.7 |
| | 17 | 11.62 | 1.02 | 9.78 | 0.72 | 0.08 | 0.02 | 8.76 | 84.1 | 6.23 | 0.69 | 0.18 | 9.8 | 13.8 | 53.7 | 54.8 | 14.1 | 25.7 | 17.3 | 640 | 4.3 |
| | 18 | 11.96 | 1.38 | 9.58 | 0.86 | 0.09 | 0.04 | 11.56 | 80.1 | 7.2 | 0.78 | 0.35 | 9.09 | 13.9 | 47.6 | 52.4 | 15.3 | 29.2 | 18.1 | 782 | 4.5 |
| | 19 | 10.44 | 0.57 | 9.15 | 0.68 | 0.02 | 0.02 | 5.5 | 87.7 | 6.52 | 0.16 | 0.15 | 9.14 | 13.4 | 49.6 | 54.3 | 14.7 | 27 | 19.5 | 643 | 4.4 |
| | 20 | 12.24 | 0.81 | 10.62 | 0.79 | 0.02 | 0 | 6.62 | 86.8 | 6.42 | 0.18 | 0.04 | 9.28 | 13.4 | 49.3 | 53.1 | 14.4 | 27.2 | 18.6 | 627 | 4.5 |
| | **Mean** | **11.45** | **0.97** | **9.666** | **0.74** | **0.06** | **0.02** | **8.41** | **84.5** | **6.46** | **0.49** | **0.18** | **9.356** | **13.5** | **50.4** | **53.8** | **14.5** | **26.92** | **18.3** | **693** | **4.48** |
| | **SD** | **0.731** | **0.3** | **0.595** | **0.09** | **0.03** | **0.01** | **2.406** | **2.97** | **0.47** | **0.3** | **0.11** | **0.289** | **0.3** | **2.36** | **1.04** | **0.55** | **1.482** | **0.84** | **77.7** | **0.15** |

*Highlighted means are to show where WT and *Sost*$^{-/-}$ are statistically different, compared by sex, using students ttest p<0.05

**Supplementary Table 3. List of Cell Surface Markers used for Flow Cytometry**

| Cell Type | Markers | | | | |
|---|---|---|---|---|---|
| Transplanted LT-HSCs | $Lin^-$ $Sca1^+$ $cKit^+$ (LSK) | $CD150^+$ | $CD48^-$ | | |
| All other LT-HSC | $Lin^-$ $Sca1^+$ $cKit^+$ (LSK) | $CD150^+$ | $CD48^-$ | $Flk2^-$ | $Flk2^-$ |
| ST-HSCs | $Lin^-$ $Sca1^+$ $cKit^+$ (LSK) | $CD150^-$ | $CD48^-$ | $Flk2^-$ | $Flk2^-$ |
| MPP2 | $Lin^-$ $Sca1^+$ $cKit^+$ (LSK) | $CD150^+$ | $CD48^+$ | $Flk2^-$ | $Flk2^-$ |
| MPP3 | $Lin^-$ $Sca1^+$ $cKit^+$ (LSK) | $CD150^-$ | $CD48^+$ | $Flk2^-$ | $Flk2^-$ |
| MPP4 | $Lin^-$ $Sca1^+$ $cKit^+$ (LSK) | $CD150^-$ | $CD48^+$ | $Flk2^+$ | $Flk2^+$ |
| T Cells | $Gr1^-$ | $CD3^+$ | $CD19^-$ | | |
| B Cells | $Gr1^-$ | $CD3^-$ | $CD19^+$ | | |
| Monocytes | $CD3^-$ | $CD11b^+$ | $Gr1^-$ | | |
| Granulocytes | $CD3^-$ | $CD11b^+$ | $Gr1^+$ | | |
| B Cell, Fraction A (BM) | $B220^+$ $CD43^+$ | $cKit^-$ | $CD19^-$ | | |
| B Cell, Fraction B-C (BM) | $B220^+$ $CD43^+$ | $cKit^-$ | $CD19^+$ | | |
| B Cell, Fraction D (BM) | $B220^+$ $CD43^-$ | $cKit^-$ | $CD19^+$ | $IgM^-$ | $IgD^-$ |
| B Cell, Fraction E (BM) | $B220^+$ $CD43^-$ | $cKit^-$ | $CD19^+$ | $IgM^+$ | $IgD^-$ |
| B Cell, Fraction F (BM) | $B220^+$ $CD43^-$ | $cKit^-$ | $CD19^+$ | $IgD^+$ | |
| B Cell (Spleen) | $B220^+$ | $CD19^+$ | | | |
| B Cell (Spleen) | $B220^+$ | $IgM^+$ | | | |
| Mesenchymal stromal cell | $Lin^-$, $CD45^-$ | $CD31^-$ | $Sca1^+$ | $CD51^+$ | |
| Osteoblasts | $Lin^-$, $CD45^-$ | $CD31^-$ | $Sca1^-$ | $CD51^+$ | |
| Endothelial cell | $Lin^-$, $CD45^-$ | $CD31^+$ | $Sca1^+$ | | |
| "Other" Bone cell | $Lin^-$, $CD45^-$ | $CD31^-$ | $Sca1^-$ | $CD51^-$ | |

**Supplemental Figure Legends**

*Supplemental Figure 1. Sclerostin-depleting antibodies change hematopoietic differentiation.*

(A) Bone marrow cellularity of HSPCs in PBS vs Scl-Ab treated mice; (B) Bone marrow cellularity of mature ; (C) Splenic cellularity of HSPCs; (D) Splenic cellularity of mature lineages; (E) Bone marrow frequency of B cell populations; (F) Splenic frequency of B cell populations; (G) Bone marrow cellularity of B cell populations; (H) Splenic cellularity of B cell populations. Mice were 8 weeks of age at start of treatment and all female. Seven mice received PBS, 7 mice received Scl-Ab antibody. p<0.05*, p<0.01**, p<0.001***, p<0.0001**** Mann-Whitney, non-parametric ttest.

*Supplemental Figure 2. Analysis of HSPCs in Sost$^{-/-}$ mice*

(A) Donor cellularity of BM HSPC progenitors in WT→*Sost*$^{-/-}$ chimeras; (B) Donor cellularity of spleen HSPC progenitors in WT→*Sost*$^{-/-}$ chimeras; (C) Expression of *Hoxb4* and *p21/Cip1* in LSKs normalized to *Gapdh*; (D) Cell numbers in controls and 5-FU treated WT and *Sost*$^{-/-}$ mice 5 days after 5FU injection, in 2 independent experiments; (E) Summary of basal (non-5FU-treated mice) cell cycle analysis in LSKs; (F) Summary of cell cycle analysis in WT and *Sost*$^{-/-}$ mice in LSKs 5 days after 5-FU injection. LSK frequencies in individual mice are shown as normalized to the mean of the WT 5-FU treated mice; (G) Representative flow cytometry plots of cell cycle analysis on gated LSKs in untreated and 5FU-treated animals. Male and female mice were 17 weeks of age; p<0.05*, p<0.01**, p<0.001***, p<0.0001**** Mann-Whitney, non-parametric ttest.

*Supplemental Figure 3. Evidence of extramedullary hematopoiesis in Sost$^{-/-}$ mice*

(A) Spleen weight in WT and *Sost*$^{-/-}$ mice. Values represent total spleen weight divided by total mouse weight; (B) Splenic pulp area, as calculated by area of indicated pulp type (white or red) divided by total pulp area stained; (C) H&E staining of spleens. Size reference is 500 µm. Dark purple round spots are white pulp and pink area is red pulp; (D) LT-HSC frequency analyzed over time in the bone marrow; (E) ST-HSC frequency analyzed over time in the bone marrow; (F) LT-HSC frequency analyzed over time in the spleen; (G) ST-HSC frequency analyzed over time in the spleen. In Figures 3D-3G, the data is representative of 4 mice per time point for WT and *Sost*$^{-/-}$ mice.

p<0.05*, p<0.01**, p<0.001***, p<0.0001**** Mann-Whitney, non-parametric ttest.

*Supplemental Figure 4. LT-HSCs from the spleen show no functional differences compared to the bone marrow*

(A) Experimental schematic of splenic LT-HSCs transplantation assay; (B) Bone marrow and splenic cellularity in chimeras; (C) Bone marrow and splenic donor chimerism; (D) Bone marrow cellularity of donor-derived HSPCs ;(E) Splenic cellularity of donor-derived HSPCs; (F) Bone marrow cellularity of donor-derived mature lineages; (G) Splenic cellularity of donor-derived mature hematopoietic lineage cells. Data shown are pooled from 2 separate experiments. Male or female donors with age ranges from 37-49 weeks were used. Recipient mice were male and female with age ranges from 11-15 weeks. Chimeras were analyzed 19 weeks post-transplantation. Donor LT-HSC cell numbers ranged from 1,000-3,000 LT-HSCs cells per mouse. $p<0.05$*, $p<0.01$**, $p<0.001$***, $p<0.0001$**** Mann-Whitney, non-parametric ttest.

*Supplemental Figure 5. Analysis of CXCL12 and SCF in bone stromal cells of Sost$^{-/-}$ mice*

(A) Representative FACS plots depicting gating strategy for collagenase-digested bones; (B) Relative CXCL12 gene expression in *Sost*$^{-/-}$ bone cells; (C) Relative SCF gene expression in *Sost*$^{-/-}$ bone cells; (D) Summary of bone cell frequencies after digest of femurs and tibiae of WT (n=14) and KO (n=26) mice. $p<0.05$*, $p<0.01$**, $p<0.001$***, $p<0.0001$**** Mann-Whitney, non-parametric ttest.

Supplemental Figure 1

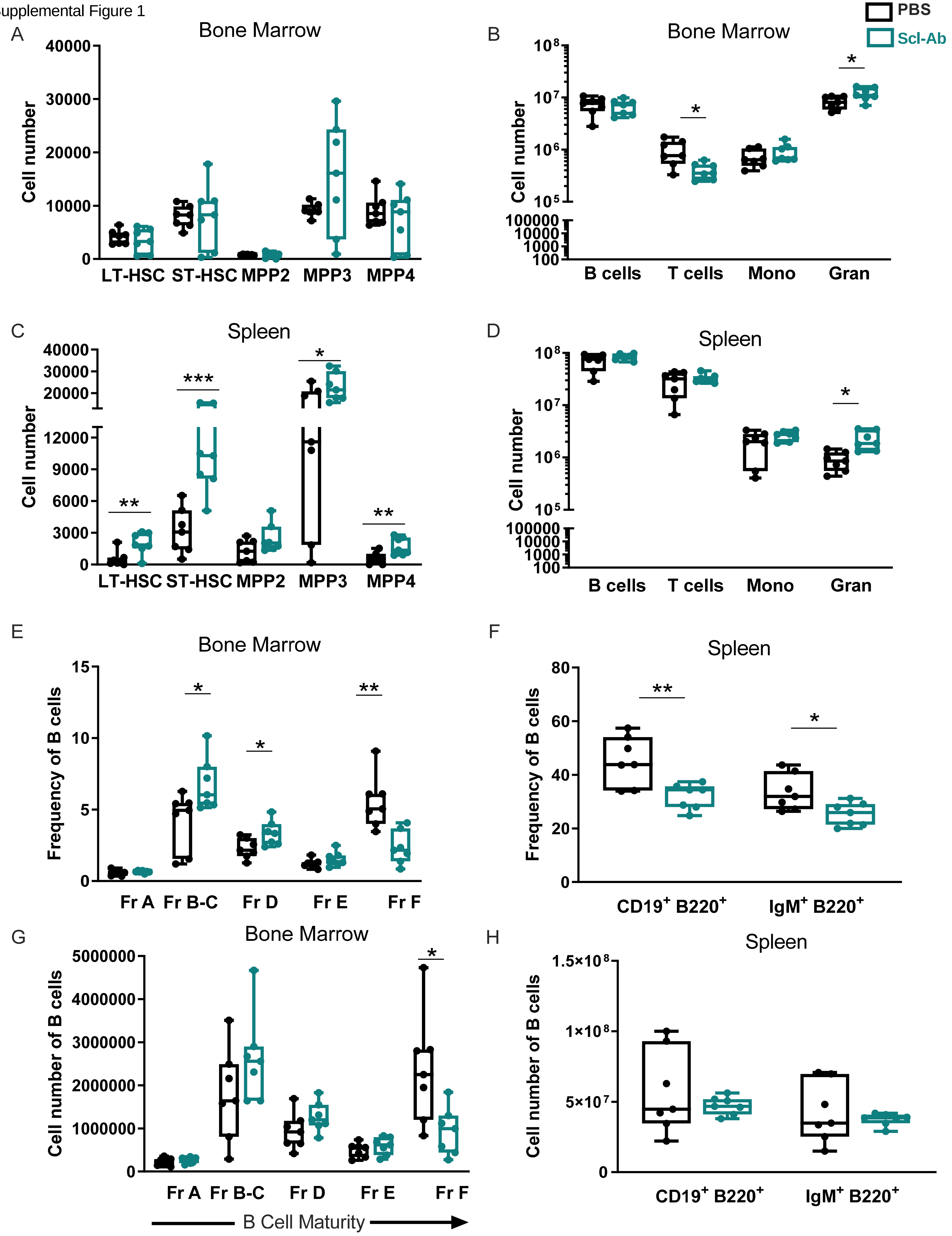

Supplemental Figure 2

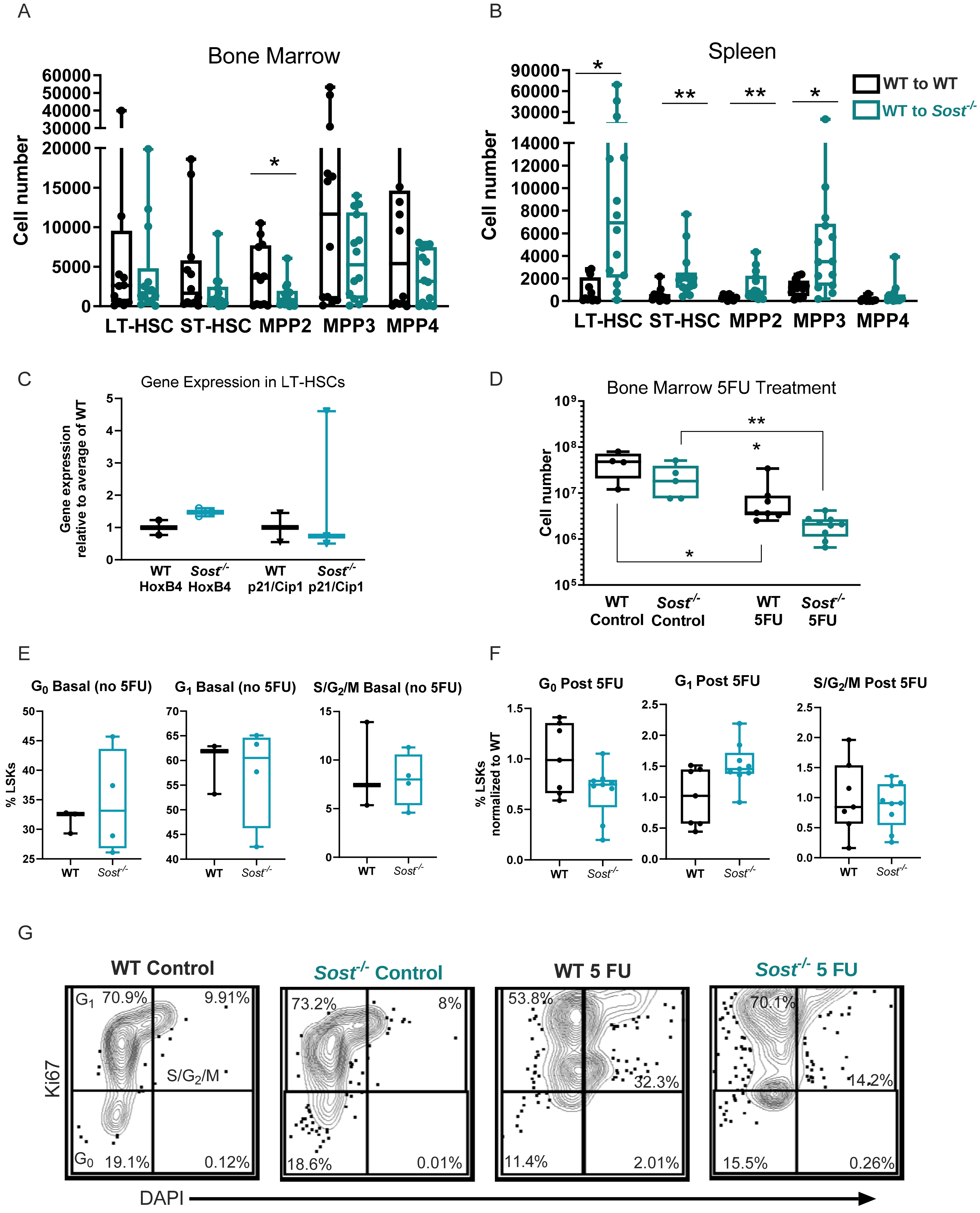

Supplemental Figure 3

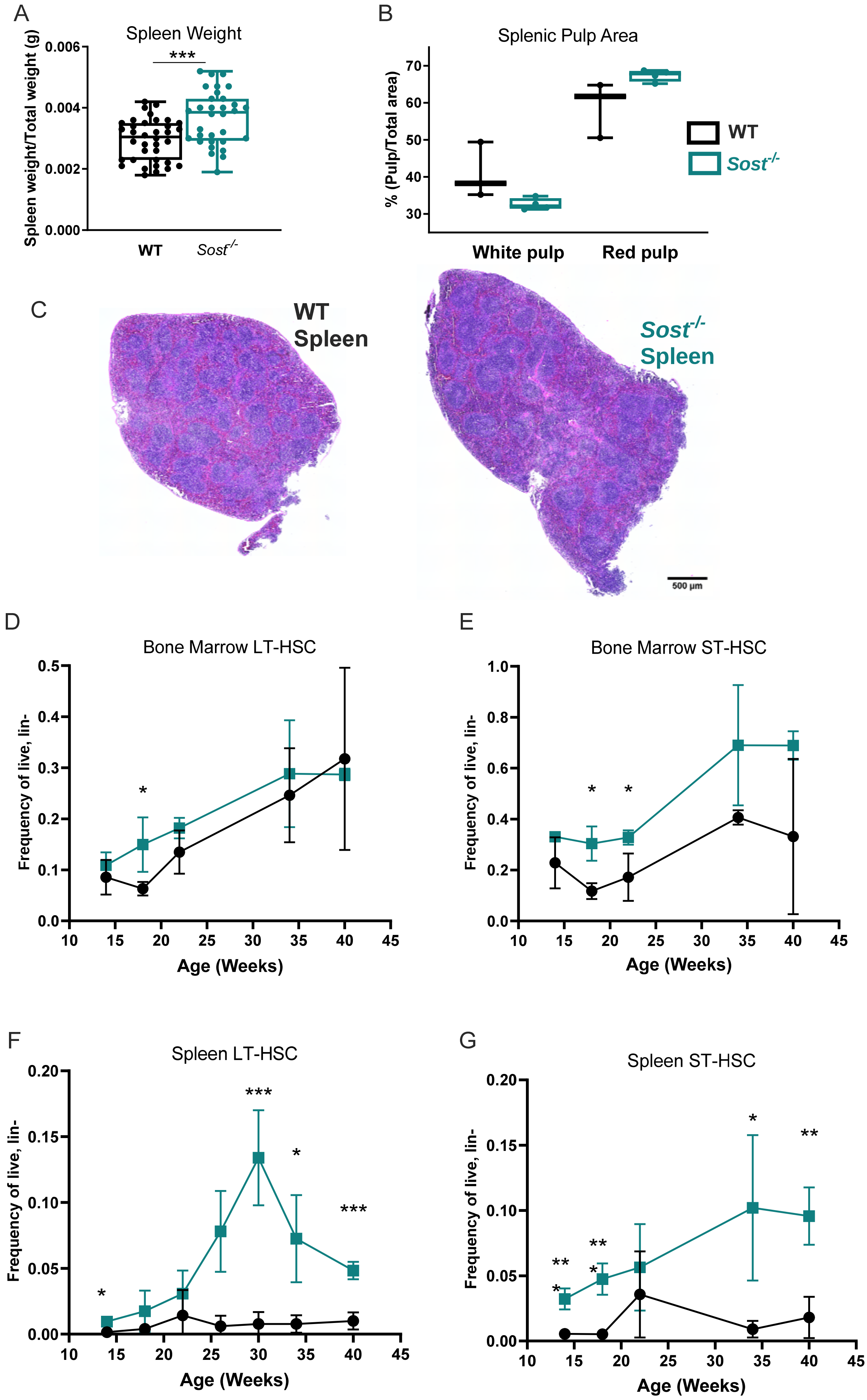

Supplemental Figure 4

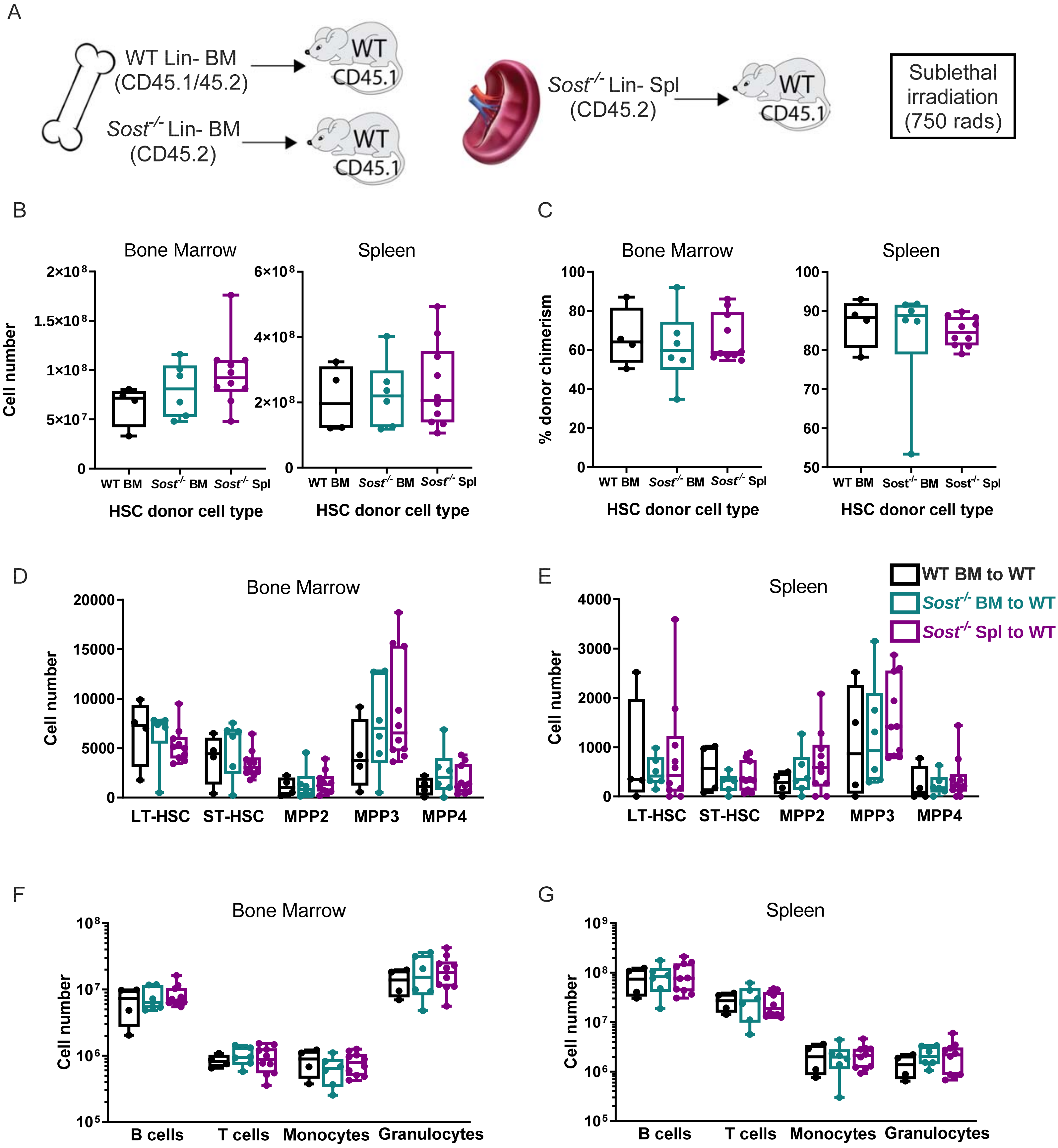

Supplemental Figure 5

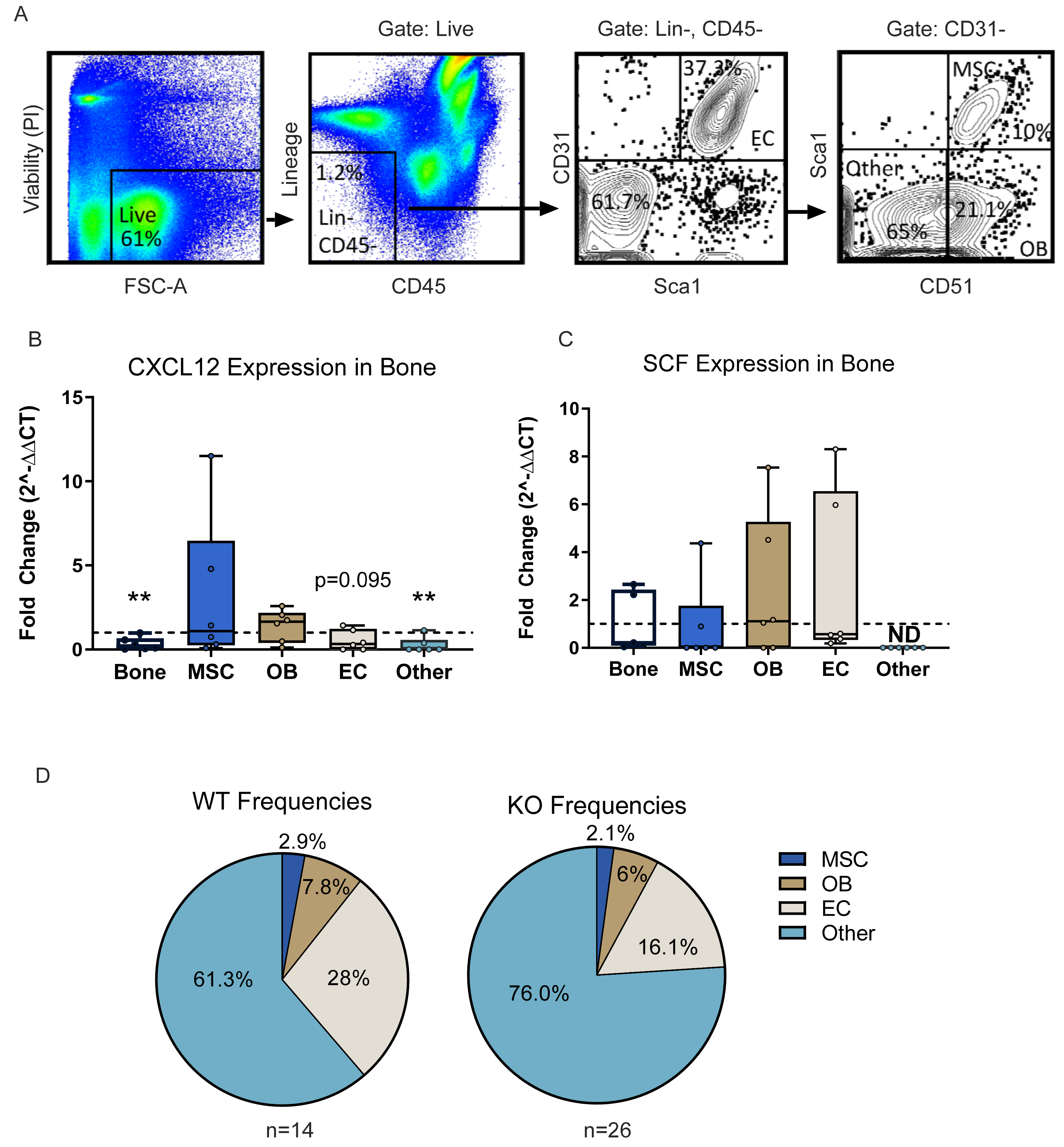
